# Using MinION nanopore sequencing to generate a *de novo* eukaryotic draft genome: preliminary physiological and genomic description of the extremophilic red alga *Galdieria sulphuraria* strain SAG 107.79

**DOI:** 10.1101/076208

**Authors:** Amanda M. Davis, Manuela Iovinella, Sally James, Thomas Robshaw, Jennifer R. Dodson, Lorenzo Herrero-Davila, James H. Clark, Maria Agapiou, Simon J. McQueen-Mason, Gabriele Pinto, Claudia Ciniglia, James P. J. Chong, Peter D. Ashton, Seth J. Davis

## Abstract

We report here the *de novo* assembly of a eukaryotic genome using only MinION nanopore DNA sequence data by examining a novel *Galdieria sulphuraria* genome: strain SAG 107.79. This extremophilic red alga was targeted for full genome sequencing as we found that it could grow on a wide variety of carbon sources and could uptake several precious and rare-earth metals, which places it as an interesting biological target for disparate industrial biotechnological uses. Phylogenetic analysis clearly places this as a species of *G. sulphuraria*. Here we additionally show that the genome assembly generated via nanopore long read data was of a high quality with regards to low total number of contiguous DNA sequences and long length of assemblies. Collectively, the MinION platform looks to rival other competing approaches for *de novo* genome acquisition with available informatics tools for assembly. The genome assembly is publically released as NCBI BioProject PRJNA330791. Further work is needed to reduce small insertion-deletion errors, relative to short-read assemblies.

## Introduction

Species in the genus *Galdieria* are unicellular red algae that exhibit wide metabolic versatility and display enormous capacity to thrive at temperatures up to 50 °C and under highly acidic conditions (Gross and Schnarrenberger 1995). These organisms are found at sulphur vents (Ciniglia, Yoon et al. 2004, Yoon, Ciniglia et al. 2006, Pinto 2007) and anthropomorphic sites contaminated with acidic conditions, such as in acid-mine drains and the River Tinto in Spain (López-Archilla, Marín et al. 2001). Isolates within this genus display a broad metabolic repertoire allowing vigorous growth on a great many sugar-based carbon sources (Gross and Schnarrenberger 1995, Qiu, Price et al. 2013). These species are metabolic workhorses. Efforts have been made to harness these abilities and develop *Galdieria* for industrial biotechnological applications (Graziani, Schiavo et al. 2013, Henkanatte-Gedera, Selvaratnam et al. 2015, Minoda, Sawada et al. 2015, Selvaratnam, Pegallapati et al. 2015, Ju, Igarashi et al. 2016). Specific goals will undoubtedly include identifying those strains best capable of using waste from the food industry and biofuel community, and assess the nature of metabolic conversions, including metal concentration bioaccumulation or bioabsorption, to determine the utility and benefit of resultant useful products. This includes an examination of the diverse *Galdieria* genomes to explore the "industrial space" of isolates of this species with an intention of creating recombinant proteins of unique benefits, as they are expected to be heat and acid tolerant.

Metabolism has been most characterised in *Galdieria sulphuraria* strain 074W. Numerous reports have shown that this organism can grow between pH levels of ~0 to about pH 5 (Gross and Schnarrenberger 1995, Gross, Küver et al. 1998, Weber, Oesterhelt et al. 2004, Schmidt, Wiebe et al. 2005, Oesterhelt, Klocke et al. 2007, Linka, Jamai et al. 2008, Qiu, Price et al. 2013). Additionally it can thrive under autotrophic growth in the light, as well as mixotrophic growth in the light. In darkness, heterotrophic growth is supported under a huge range of available carbon sources, including essentially all tested sugars, sugar-alcohols and organic acids (Gross and Schnarrenberger 1995, Qiu, Price et al. 2013). These are often metabolised, as for example, *G. sulphuraria* can thrive in the presence of nearly 1M sucrose and 0.2 M ammonium sulphate without any alterations in cell division rates and general growth (Schmidt, Wiebe et al. 2005). It is also tolerant to massive concentrations of cations, including diverse heavy metals, as well as anions (Dopson and Holmes 2014). Additionally, perhaps as acidic environments increase the solubility of most cations, *G. sulphuraria* bioaccumulates numerous rare-earth metals and several precious metals (Minoda, Sawada et al. 2015, Ju, Igarashi et al. 2016). In this versatile growth state, interest in this organism has been seen in the industrial biotechnology sector, both for remediation of wastes and in the production of high value chemicals (Schmidt, Wiebe et al. 2005, Selvaratnam, Pegallapati et al. 2014, Henkanatte-Gedera, Selvaratnam et al. 2015, Selvaratnam, Pegallapati et al. 2015, Henkanatte-Gedera, Selvaratnam et al. 2016).

*Galdieria* genomics emerged with the deep sequencing of an Expressed Sequence Tags (EST) library from the type species *Galdieria sulphuraria* 074. It was estimated that sequence reads covered >70% of the *G. sulphuraria* genome (Weber, Oesterhelt et al. 2004). These were compared to the thermo-acidophilic alga *Cyanidioschyzon merolae*, which is believed to be autotrophic (Barbier, Oesterhelt et al. 2005). Consistent with versatile hetero- and mixo-trophic grown on most carbon sources tested, within the subset of genes unique to *Galdieria*, many were found to encode metabolite transporters and enzymes (Barbier, Oesterhelt et al. 2005). This supports the notion of a broad metabolic repertoire within *Galdieria* that is absent from related organisms in the Family *Cyanidiaceae* (Schönknecht, Chen et al. 2013, Schönknecht, Weber et al. 2014). Building on this, *Galdieria* genomes have been reported based on traditional and second-generation approaches.

Here we generated a draft eukaryotic genome using the MinION third-generation nanopore sequencing method. The selected strain of *G. sulphuraria* was chosen as it is available at several stock centres and performed well under numerous lab-based tests. This 107.79 strain was first selected over other available lines as it notably grew quickly on agar plates resulting in larger colonies than other tested *Galdieria* species. Here we found that 107.79 could grow under massive amounts of diverse anions and could bioaccumulate several rare-earth and precious metals. It also could grow heterotrophically on diverse carbon sources, but not all of those that support growth in *G. sulphuraria*. Waste sources supported vigorous growth implying resistance to toxicity of various metals. In considering its genome, we strive to compare Illumina paired-end runs to MinION runs alone, or read corrected with Illumina reads. Together we found that long-sequence reads could generate a nearly contiguous genome, and that *Galdieria* sp. 107.79 is an excellent candidate for further bioinformatics and phycological analyses.

## Materials and methods

The studied *Galdieria* species SAG 107.79 was obtained from Culture Collection of Algae at the University of Göttingen, Germany (SAG). It is also available at The Culture Collection of Algae at the University of Texas at Austin (UTEX) as strain UTEX 2393 and the algal collection at the Dipartimento di Biologia, Università Federico II of Naples (ACUF) as strain ACUF 139. It was previously erroneously classified as *Cyanidium caldarium* (Merola, Castaldo et al. 1981, Moreira, López-Archilla et al. 1994).

Our standard media for growth was 4.4 g / L Murashige and Skoog + vitamins (Duchefa M0222) supplemented with 10 g / L sucrose. The pH was adjusted to 2.0 with sulphuric acid and was sterilized by heat. Sugars and other carbon tests were from chemicals acquired from Sigma or Amazon.uk. In some experiments, pH was adjusted by a metalliferous waste acid that was predominantly phosphoric acid (obtained from Airdale, UK). Rare earth and precious metal salts were all from Sigma, as were concentrated acids. The acid recoverable solution "Cyanidium Medium" was as described at the SAG stock centre (Koch 1964), from a soil sample taken from the Walmgate Stray (53.9467°N 1.0622°W). Briefly, to a soil sample, two parts water was added and sulphuric acid was added until the pH became 1.4. This solution was filtered for particles over Whatman paper and the resultant supernatant was heat sterilised.

Microscopy entailed imaging stationary cultures grown autotrophically on a Zeiss LSM 710 meta upright.

Carbon growth tests have been performed were growth was detected in the dark on 10 g/L for: sucrose, glucose, arabinose, fructose, galactose, maltose, lactose, glycerol, erythritol, xylitol, manitol, sorbitol, isomalt, myoinositol, thiamine, biotin, adenine, glycine and di-glycine.

For the high anions comparison, media were prepared with the addition of 1% (v/v) of each of the following concentrated acids: HCl, H_2_SO_4_, HNO_3_ and H_3_PO_4_. The specific concentrations of the stock acids available were taken into account and added volumes were adjusted accordingly. The pH of all media was increased to 2 using KOH. Cultures (25mL) were incubated at 38°C for 19 days, in pre-weighed plastic centrifuge tubes. The cultures were given airings, under aseptic conditions, every few days, to encourage culture growth. The dried cell fraction was attained by pelleting the cultures by centrifugation, decanting the supernatant, then washing with deionised water, centrifuging again, washing again and centrifuging for a final time. The water was decanted and the centrifuge tubes weighed to calculate the fresh algal weight. The cultures were placed in an air-drying oven for 18 hrs at 60°C, then weighed again to calculate the dry algal weight.

To assess growth under a synthetic condition, a media was formulated entitled <u>Acid Mine Drainage Emulating Media</u> (Heavy Metals Uptake Comparison). This medium was prepared as our standard MS-sucrose pH2, with the following additions: 2mM FeNaEDTA, 1.5mM CuCl_2_, 0.2mM NiCl_2_.6H_2_O, 0.2mM CoCl_2_.6H_2_O. Metals were added after autoclaving, to prevent any premature precipitation. Cultures (40 mL) were incubated at 38°C for 7 days, in sterile conical flasks, transferred to pre-weighed centrifuge tubes for accurate weight determinations. The cells were dried and weighed as per the high-level anions comparison. The dried cell fractions were submitted for ICP-MS analysis, as described (Parker, Rylott et al. 2014), along with a sample of the original medium (filtered), to allow approximate percentage recoveries to be calculated.

For a Precious Metals uptake comparison, the standard MS-sucrose medium was supplemented with 1 × 10^−3^mM PdCl_2_, 1 × 10^−3^mM AgNO_3_, 5 × 10^−4^mM PtCl_2_ and 5 × 10^−4^mM AuBr_3_. Pd, Pt and Au were first dissolved in concentrated HCl. The solution was adjusted to a pH of ~2 with NaOH, then diluted with water to make a stock solution 1000 times more concentrated than the final concentrations required. This was 0.45μM filter sterilised, then added to the medium after autoclaving. As Ag is known to precipitate in the presence of chloride, AgNO_3_ was added to the individual centrifuge tubes immediately before the addition of parent culture, to ensure the amount of Ag was equal for each culture. Cultures (25 mL) were incubated at 38°C in pre-weighed centrifuge tubes, laid horizontally, for 8 days. They were then transferred to sterile conical flasks, and incubated for a further 8 days, The cells were dried and weighed as per the high-level anions comparison. The dried cell fractions were submitted for ICP-MS analysis, along with a sample of the original medium.

For a Lanthanides uptake comparison, the MS-sucrose medium was supplemented with 6 × 10^−4^mM Y(NO_3_)_3_.6H_2_O, 7 × 10^−4^mM LaCl_3_, 6 × 10^−4^mM GdC_3_F_9_O_9_S_3_, 7 × 10^−4^mM NdF_3_ and 3 × 10^−4^mM YbC_3_F_9_O_9_S_3_. All elements were dissolved in water, then diluted with further water to make a stock solution 1000 times more concentrated than the final concentrations required. The pH was adjusted to ~2 with H_2_SO_4_ before the dilution was completed. Cultures (25mL) were incubated at 38°C in pre-weighed centrifuge tubes, laid horizontally, for 8 days. They were then transferred to sterile conical flasks and incubated for a further 8 days. The cells were dried and weighed as per the high-level anions comparison. The dried cell fractions were then submitted for ICP-MS analysis.

For phylogenetic analysis of strain 107.79, this was performed relative to 37 taxa belonging to different genera and species of Cyanidiophyceae (2 *Cyanidioschyzon merolae*, 3 *Cyanidium caldarium*, 2 mesophilic *Cyanidium*, 6 *G. phlegrea*, 10 *G. maxima*, 12 *G. sulphuraria*, 1 *G. daedala*, 1 *G. partita*) and belonging to the Algal Culture Collection of University Federico II of Naples (ACUF www.acuf.net), were used for the traditional Sanger-sequencing method, the strain *G. sulphuraria* 107.79, belonging to the UTEX Algal Culture Collection was used for *MinION sequencing*. For DNA extraction, algal cells were ground with glass beads using a Mini-BeadBeater (BioSpec, Bartlesville, Oklahoma, USA) operated at 13,000 rpm per min for 5 min. Total genomic DNA was extracted using the DNeasy Plant Mini Kit (Qiagen, Santa Clarita, California, USA). PCR and sequencing reactions were conducted using specific primers for three plastid genes, psbA, psaA and rbcL (Ciniglia, Yoon et al. 2004, Ciniglia, Yang et al. 2014). The polymerase chain reaction (PCR) products were purified with the QIAquick PCR purification kit (Qiagen) and used for direct sequencing using the BigDyeTM Terminator Cycle Sequencing Kit 3.1 (PE-Applied Biosystems, Norwalk, Connecticut, USA) and an ABI-3500 XL at the Microgem Laboratory (Naples, Italy). Forward and reverse electropherograms were assembled and edited using the program Chromas Lite v.2.1 (www.technelsium.com.au/chromas.html). New sequences were aligned with already published sequence data (Ciniglia, Yoon et al. 2004, Toplin, Norris et al. 2008, Skorupa, Reeb et al. 2013, Ciniglia, Yang et al. 2014, Hsieh, Zhan et al. 2015), obtained from GenBank (www.ncbi.nlm.nih.gov), using BioEdit Sequence Alignment Editor (http://www.mbio.ncsu.edu/BioEdit/bioedit.html). No gaps or indels have been incorporated in the alignments. The dataset used here contained 5 outgroup and 37 taxa, comprising a total of 2965 DNA positions (1080 bp for rbcL, 899 bp for psbA and 980 for psaA).

Phylogenetic trees were inferred with maximum likelihood (ML) and Bayesian inference methods. Maximum likelihood (ML) analyses were performed using the TOPALi v2.5 (Milne, Lindner et al. 2009). The best likelihood tree was estimated under the general time reversible (GTR) substitution with gamma distributed rate heterogeneity (G) model. One thousand bootstrap analyses (MLB) were performed using the same program option. The Bayesian inference was obtained through the algorithm of MrBayes implemented in Topali v2.5 (Milne, Lindner et al. 2009). Two runs were made with 1 million generations and a sample frequency of 200 using the same model for the Maximum Likelihood analyses (GTR + G + I). The first 25% of sampled trees were discarded as burnin before calculating posterior probabilities. Consensus tree topology and its log-likelihood mean (lnL) were also determined for each dataset as well as substitution model parameters as the 95 % credibility interval for the gamma distribution shape (α).

### MySeq Illumina data

For DNA extraction used for Illumina, DNA was extracted by resuspending a stationary phase algal paste with DNA extraction buffer (Davis, Hall et al. 2009). This was incubated for 1 hr at 65°C, centrifuged and the supernatant was precipitated by the addition of 1: 1 isopropanol. A resultant pellet was suspended in Qiagen buffer PB, applied to a miniprep column and washed according to manufacturers' details. DNA was eluted by applying EB to the column where 200mL of 65 °C EB was added in 4 sequential elution steps. Sequencing was as described (Willing, Rawat et al. 2015). After trimming, Illumina MiSeq reads were assembled using Spades v3.1 (Bankevich, Nurk et al. 2012).

### MinION sequencing

A DNA library was prepared by shearing DNA in a Covaris, and then generating a library with a Oxford Nanopore SQK-006 library preparation kit using the standard ONT protocol (Oxford Nanopore Technologies), as per manufacturers' instructions. This was then run on a R7.3 flow cell (ONT) for 36 hours. After running on the MinION, base calling was performed using Metrichor (Jain, Fiddes et al. 2015). Those reads that passed 2-D base calling were then run through nanocorrect (Loman, Quick et al. 2015), assembled using minimap and miniasm (Li 2016). The consensus sequence from the raw contigs were then improved by processing twice with monopolist (Goodwin, Gurtowski et al. 2015, Loman, Quick et al. 2015). The genome assembly is publically released as BioProject PRJNA330791. This is currently found at http://www.ncbi.nlm.nih.gov/bioproject/330791.

## Results

### General growth

The growth of *G. sulphuraria* 107.79 in MS media resulted in cells that were 5 to 10 μM in size (Fig 1a). Fluorescence overlay on an FITC channel revealed clear plasid composition (Fig 1b). On agar plates, the strain generated ~10 mm diameter-sized colonies after ~7 days growth at 37°C, and notably, mixotrophic growth had the best growth rate on plates (Fig 1c). Heterotrophic cultures were a brownish-yellow colour, and photoautotrophic cultures where sucrose was omitted resulted in a green colour. In assessing the tolerance of strain 107.79 to anions, different acid sources were compared. Whereas it is known that *G. sulphuraria* is tolerant to sulphuric acid, we had no expectations of growth on other inorganic acid sources. In comparing final growth levels of 1% nitric, hydrochloric and phosphoric acid, to 1% sulphuric acid, this strain grew equivalently under all tests. Similar fresh weight and dry weights were seen (Fig 1d). Collectively *G. sulphuraria* 107.79 appears to be broadly tolerant to numerous acid sources.

**Figure 1.**
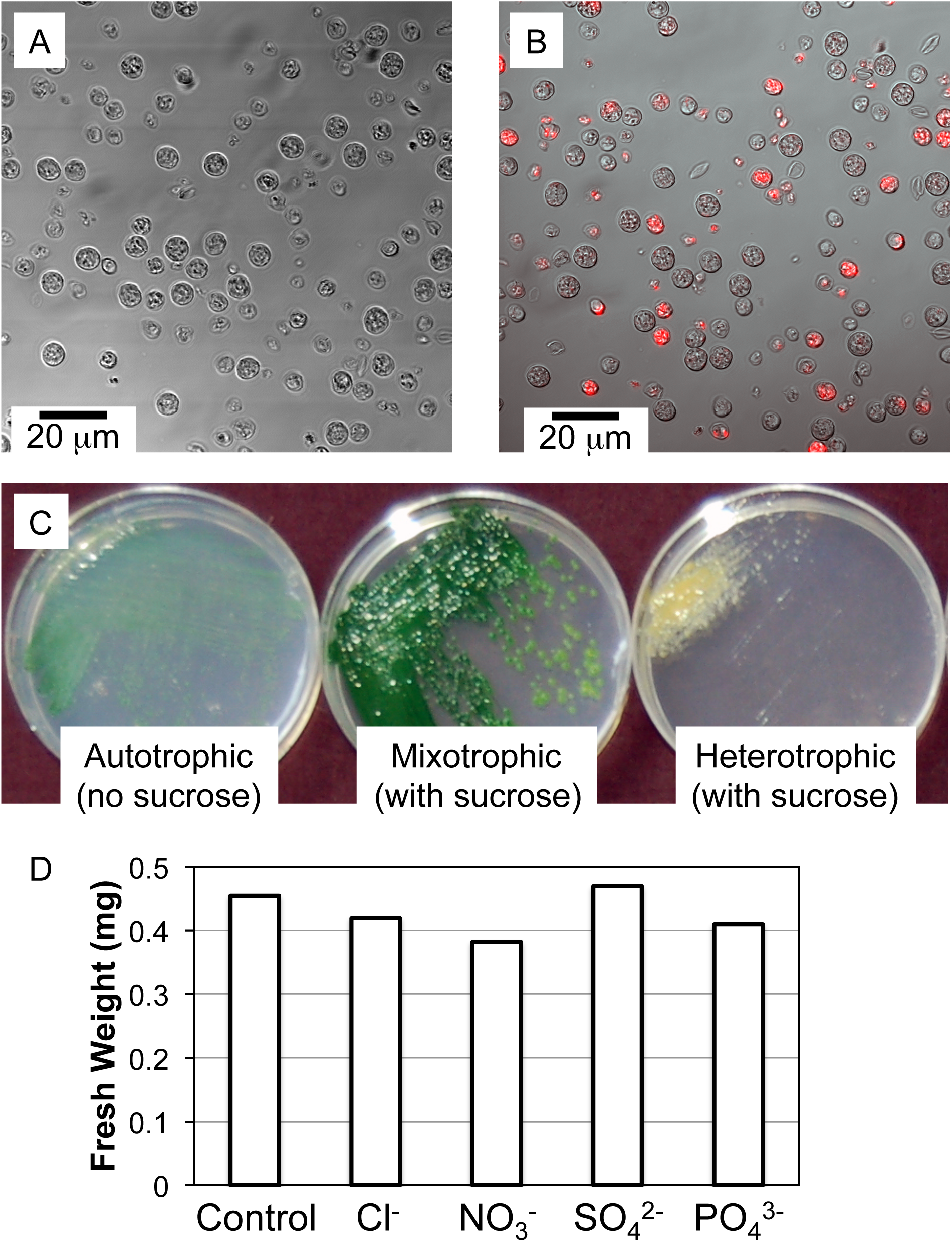
Galdieria sulphuraria SAG 107.79 growth. (A) Microscopic image after 2- weeks growth in liquid medium. (B) A separate microscopic image with confocal FITC filter set fluorescence image overlaid. (C) Plate growth after 2 weeks. (D) Dense liquid culture growth in the presence of high concentrations of various anions.

### Growth under various ionic conditions

Next *G. sulphuraria* 107.79 was tested for growth under conditions typically found at anthropomorphic acid-mine tailings. Under the presence of high Fe, Co, Cu and Ni, this strain grew well. After harvesting cells, whether metals were uptaken was assessed by ICP analysis. Fe and Ni appeared to be more accumulated than Cu and Co, but all 4 could be readily detected in the bioaccumulated fraction (Fig 2a).

**Figure 2.**
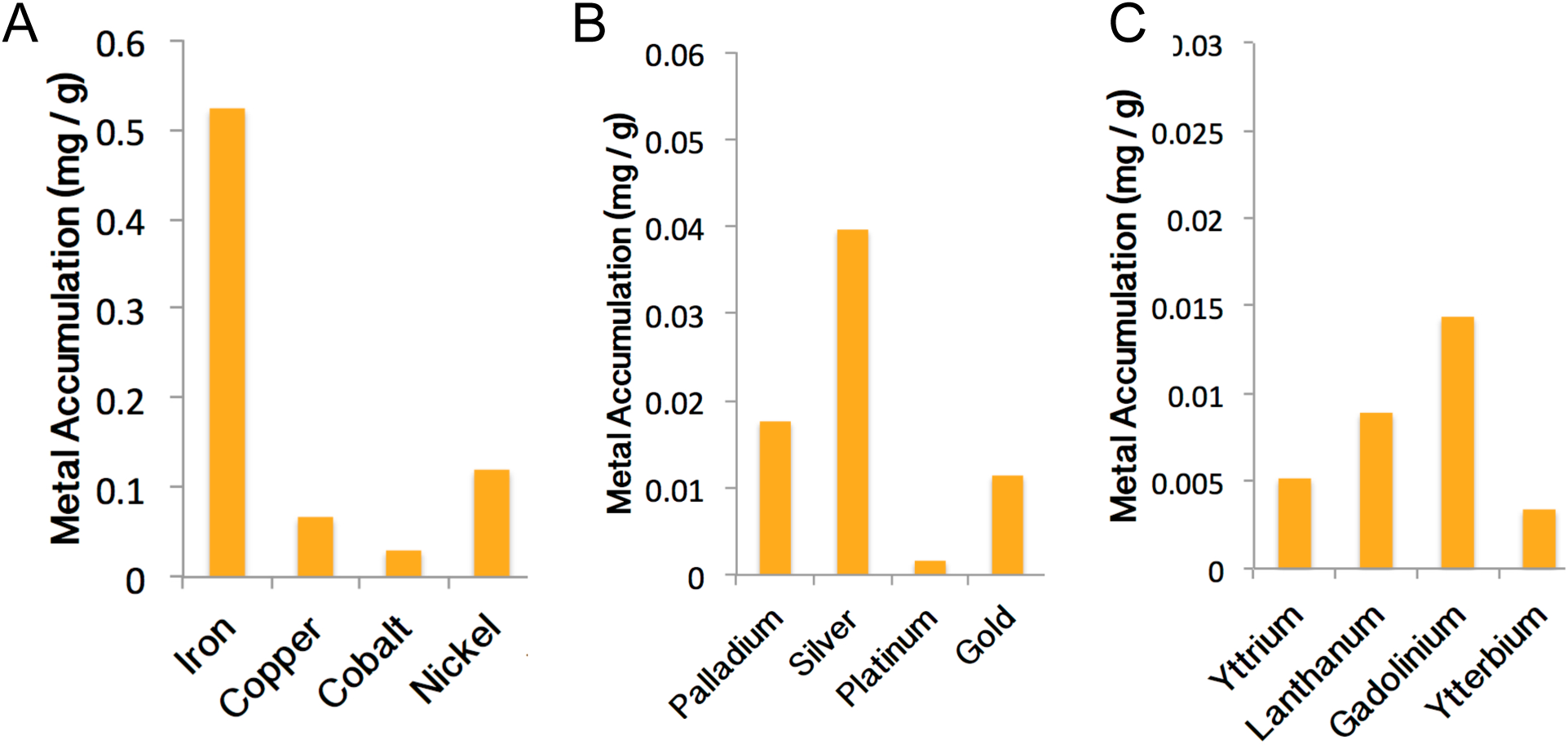
Metal uptake by strain 107.79. (A) Metal uptake after growth on acid mine tailings. (B) Precious metals uptake. (C) Rare-earths uptake.

It has been shown that *G. sulphuraria* strain 074W can take up various precious and rare-earth metals (Ju, Igarashi et al. 2016), so we assessed 107.79's ability to similarly bioaccumulate these. In a media mixture of 4 precious metals, high levels of Ag were taken up, and excellent recovery of Pd and Au was noted (Fig 2b). Like strain 074W, 107.79 does not seem to take in Pt. Similarly, in a mixture of Lanthanides, strain 107.79 appeared to bioaccumulate many of these elements (Fig 2c) in a manner seen with 074W (Minoda, Sawada et al. 2015). Together, strains 074W and 107.79 seem similar in their ability to bioaccumulate metals of high economic value from dilute media.

### Phylogenetic position of *G. sulphuraria* 107.79

We next sought to (re)establish the phylogenetic position of strain 107.79, to confirm its placement. Phylogenetic analyses based on the concatenated dataset of the three plastidial genes (rbcL, psaA and psaA), along with the new sequences generated by the MinION technology for the strain SAG107.79, revealed the clustering of this strain together with *Galdieria sulphuraria* SAG108.79 with the support of a high value of bootstrap (99%) and a Bayesian posterior probability of 1 (Fig 3). The entire cluster group together with Icelandic strains (ACUF 424, ACUF 388, ACUF 416, ACUF 402, ACUF 399, ACUF 461 and ACUF 453), along with *Galdieria partita* IPPAS P500 and *Galdieria daedala* IPPAS P508 (MLB=62%, posterior probability = 0.97). On the other branch of *Galdieria sulphuraria* clade, Turkish and Italian strains (ACUF 768, ACUF 011 and ACUF 018) took place (MLB = 100%, posterior probability =1). The phylogeny for the strain SAG107.79, using the plastid genes from MinION, confirmed the phylogeny highlighted in (Ciniglia, Yoon et al. 2004), where the 107.79 sequence used came from the Sanger method.

**Figure 3.**
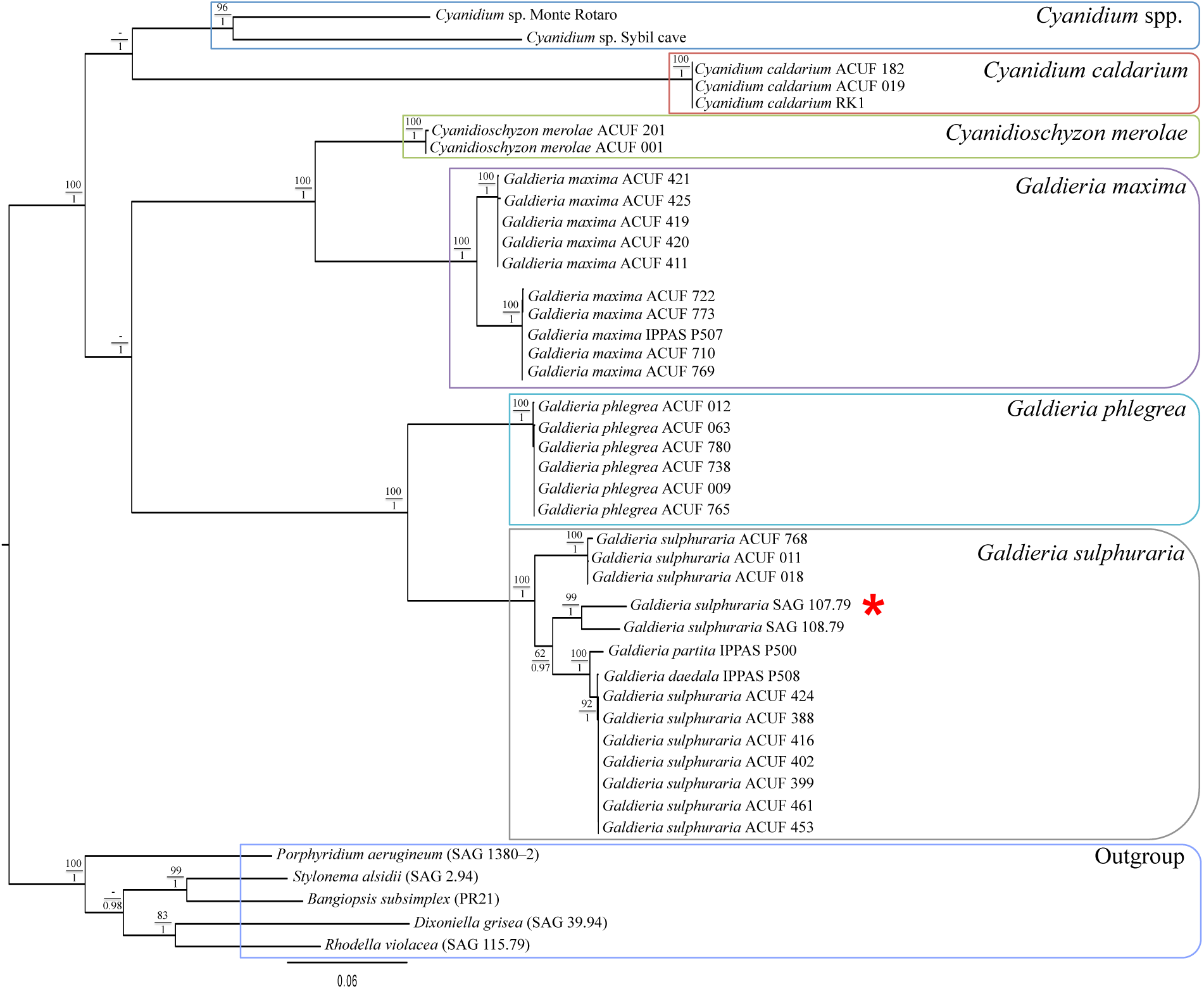
Maximum likelihood tree of *Cyanidiophyceae* inferred from RAxML analysis based on the combined plastid DNA sequences of psaA, psbA, and rbcL. Bootstrap support values are indicated near nodes above the branches, whereas the Bayesian posterior probability for clades using the GTR model is shown below the branches. Starred red is the position of the SAG 107.79. Only bootstrap values > 60% are shown.

### Generating a genome with nanopore technology

We used Oxford Nanopore technologies (ONT) MinION nanopore sequencing to define the genome of this *G. sulphuraria* species. Using a single flow cell, we generated 17,800 reads from individual DNA molecules in 2.5 hours with a modal read length of 15 kb (equivalent to 7.5 kb double stranded). Over 36 hours, this run generated a total of 1.2 Gb of sequence. The longest double-stranded read from a single molecule was 116.7 kb. In parallel, we sequenced the same organism using Illumina MiSeq technology. Together we could use this data to separately assemble de novo genomes, and a preliminary overview is now described of the results.

To examine the potential of the sequence we had generated, we used minimap/miniasm (Li 2016) to *de novo* assemble minION data. Assembly took less than three minutes (on a standard computing cluster) and produced a total assembly size of 12,092,737 bp in 127 contigs. The largest contig was 319,606 bp. By way of comparison, the short read data was assembled using Spades v3.1 assembler generated a total assembly size of 20.5 Mb, in 12,708 contigs with a largest contig of 150,370 bp (Table 1).

We have started a visual inspection to establish sequence error rates of the 107.79 genome from Illumina versus MinION data. As a rough comparison of sequence accuracy, we isolated a contig containing the Galdieria Heme oxygenase [Notated as Gasu_19790 in the reference genome; (Qiu, Price et al. 2013)], and performed an alignment. As seen (Fig 5), several sequence insertions and deletions (indels) resulted. Future Sanger sequencing is proposed to clarify whether the Illumina versus MinION data quality was the source of the sequence alignment incongruity.

## Discussion

*Galdieria* is an acidophilic and thermophilic red alga that holds enormous promise as an industrial biotechnological resource. Examining its capacity to convert acid-rich waste carbon to higher-value chemicals, we have started to examine the broad applications of bespoke industrial utility. Methods for engineering *Galdieria* start from a genomic examination of "available" enzymes. Defining multiple Galdieria genomes from diverse biogeographic habitats would support that discovery. We targeted the *G. sulphuraria* 107.79 genome as it was reported to show robust growth characteristics (Albertano, Ciniglia et al. 2000, Cozzolino, Caputo et al. 2000), which we could confirm (Figs 1-2).

One of the major challenges associated with short-read 2^nd^-generation DNA sequencing technologies is the *de novo* assembly of large contigs. In the absence of scaffolding information such as a pre-existing reference genome this problem increases with the size and complexity of the genome (or metagenome) being studied. One approach to tackling this problem is to generate long reads using solutions such as those provided by Pacific Biosciences (Faino, Seidl et al. 2015) and Oxford Nanopore Technologies (ONT) (Goodwin, Gurtowski et al. 2015). While the accuracy of these reads is lower than that achieved by short read technologies, it has been established that mapping short reads to a Pacbio scaffold has the potential to substantially increase the size of contigs that can be assembled. (Koren, Schatz et al. 2012). To start to evaluate this with MinION data, we generated a genome using nanopore sequencing (Fig 4).

**Figure 4.**
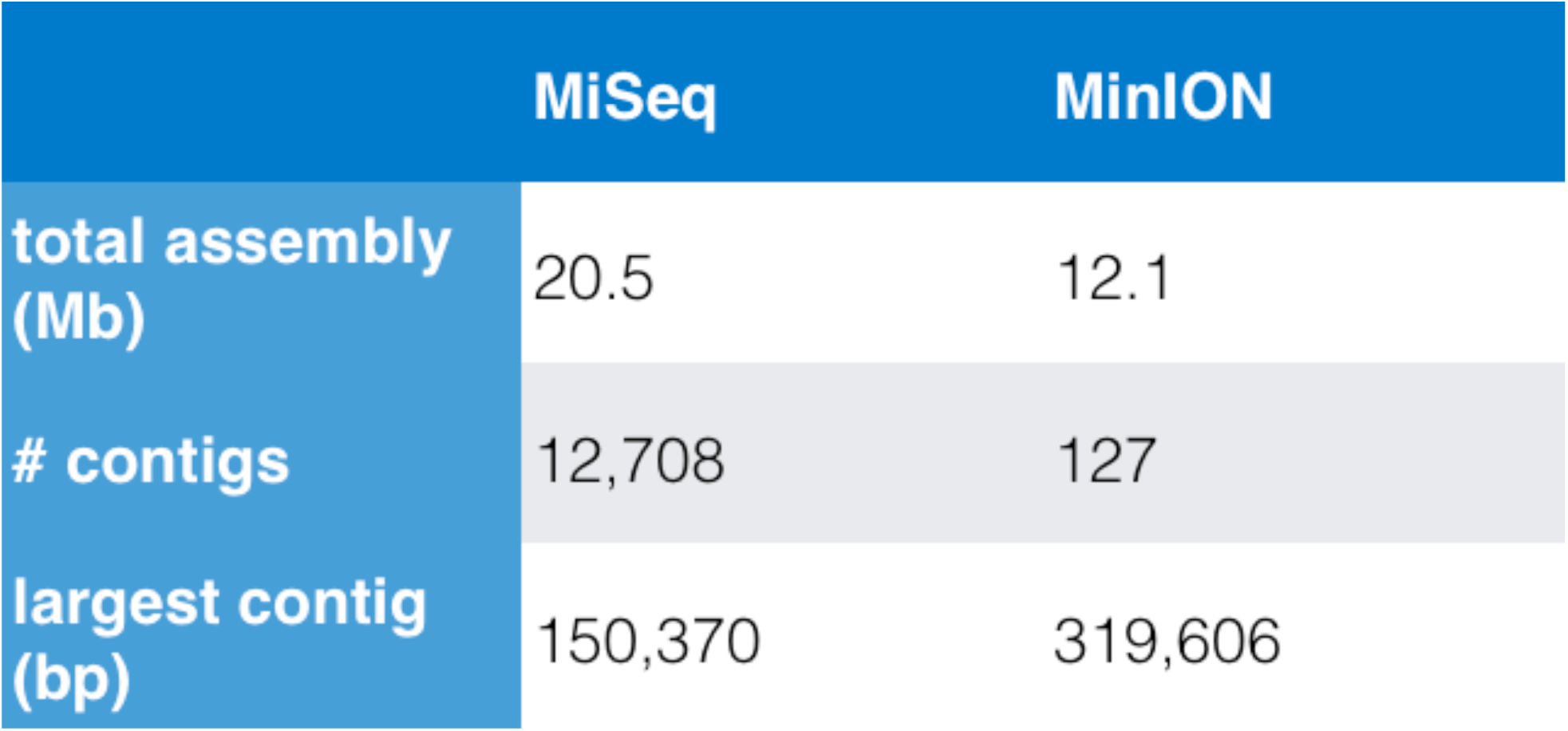
Preliminary assembly statistics of Illumina data and MinION data

The data generated by MinION sequencing exceeded our expectations. In addition to having the potential to act as a scaffold for more accurate short read assembly, we have been able to use the MinION data alone to rapidly assemble *de novo* an accurate and essentially complete genome, and we have used information from this data to position strain 107.79 within a phylogeny, concluding its expected position (Fig 3). Our results demonstrate that with suitable post-acquisition processing, long reads alone can be used to produce accurate assemblies, although these need to be improved (Fig 5). ONT's MinION technology appears to offer a number of potential advantages to provide rapid, affordable NGS capacity for *de novo* sequencing projects. These include: very low capital outlay for equipment, low input DNA requirements and simple, rapid library preparation. To the best of our knowledge this is the first report of the *de novo* assembly of a eukaryotic genome using MinION reads alone. Thus, nanopore sequencing technology and the assembly pipelines developing around this technology have allowed us to close a major gap between our ability to sequence DNA from unknown organisms and to informatically interpret these results. The low DNA sample requirements, and costs of obtaining sequence add to the attractiveness of this technology.

**Figure 5.**
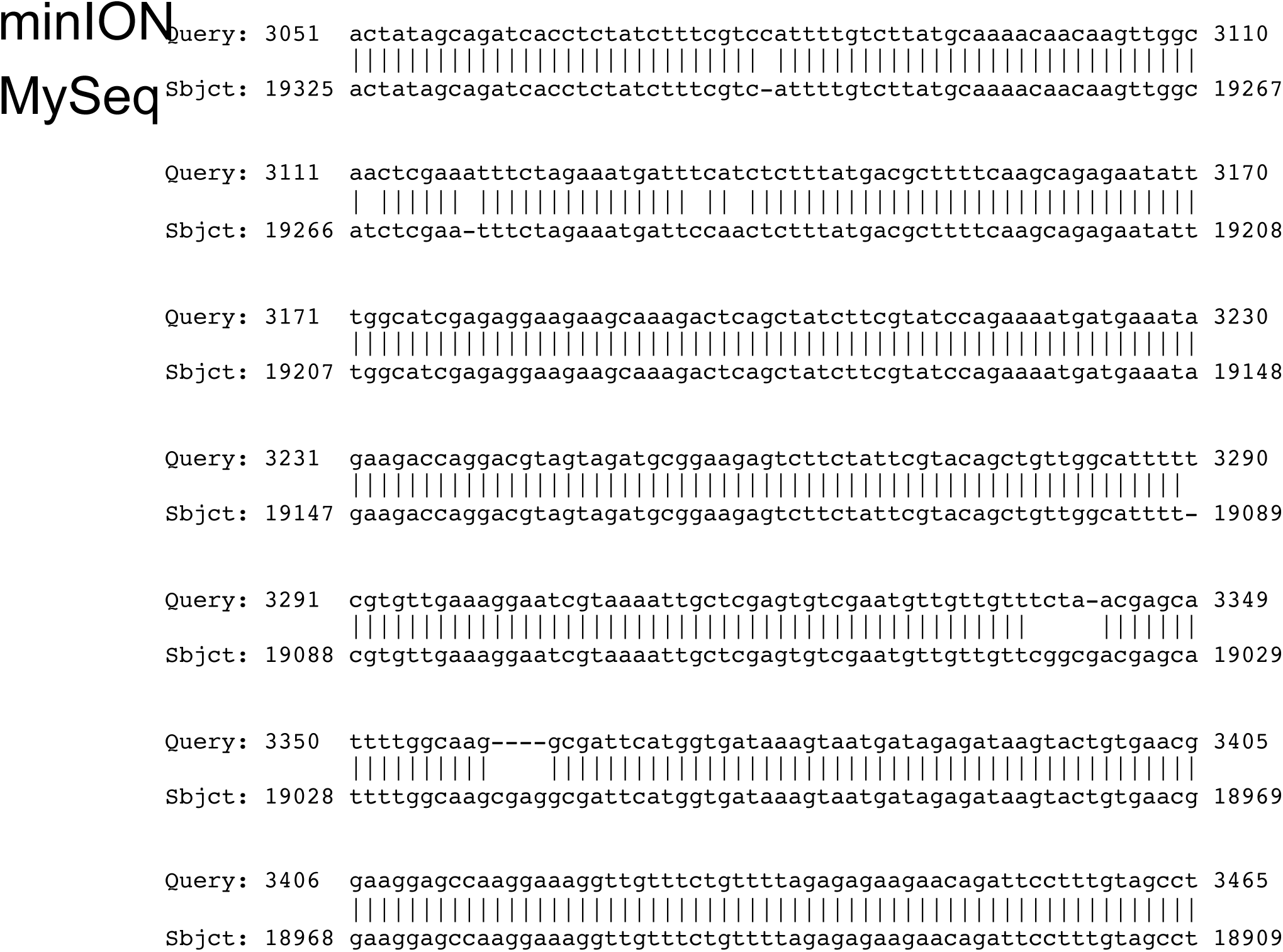
Alignment of the heme oxygenase gene of 107.79 from MinION and Illumina reads.

## References

Albertano, P., C. Ciniglia, G. Pinto and A. Pollio (2000). "The taxonomic position of Cyanidium, Cyanidioschyzon and Galdieria: an update." Hydrobiologia 433(1–3): 137–143.

Bankevich, A., S. Nurk, D. Antipov, A. A. Gurevich, M. Dvorkin, A. S. Kulikov, V. M. Lesin, S. I. Nikolenko, S. Pham and A. D. Prjibelski (2012). "SPAdes: a new genome assembly algorithm and its applications to single-cell sequencing." Journal of Computational Biology 19(5): 455–477.

Barbier, G., C. Oesterhelt, M. D. Larson, R. G. Halgren, C. Wilkerson, R. M. Garavito, C. Benning and A. P. Weber (2005). “Comparative genomics of two closely related unicellular thermo-acidophilic red algae, Galdieria sulphuraria and Cyanidioschyzon merolae, reveals the molecular basis of the metabolic flexibility of Galdieria sulphuraria and significant differences in carbohydrate metabolism of both algae." Plant physiology 137(2): 460–474.

Ciniglia, C., E. C. Yang, A. Pollio, G. Pinto, M. Iovinella, L. Vitale and H. S. Yoon (2014). “Cyanidiophyceae in Iceland: plastid rbc L gene elucidates origin and dispersal of extremophilic Galdieria sulphuraria and G. maxima (Galdieriaceae, Rhodophyta).” Phycologia 53(6): 542–551.

Ciniglia, C., H. S. Yoon, A. Pollio, G. Pinto and D. Bhattacharya (2004). "Hidden biodiversity of the extremophilic Cyanidiales red algae." Molecular Ecology 13(7): 1827–1838.

Cozzolino, S., P. Caputo, O. De Castro, A. Moretti and G. Pinto (2000). "Molecular variation in Galdieria sulphuraria (Galdieri) Merola and its bearing on taxonomy." Hydrobiologia 433(1–3): 145–151.

Davis, A. M., A. Hall, A. J. Millar, C. Darrah and S. J. Davis (2009). "Protocol: Streamlined sub-protocols for floral-dip transformation and selection of transformants in Arabidopsis thaliana." Plant Methods 5(1): 3.

Dopson, M. and D. S. Holmes (2014). "Metal resistance in acidophilic microorganisms and its significance for biotechnologies." Applied microbiology and biotechnology 98(19): 8133–8144.

Faino, L., M. F. Seidl, E. Datema, G. C. van den Berg, A. Janssen, A. H. Wittenberg and B. P. Thomma (2015). "Single-molecule real-time sequencing combined with optical mapping yields completely finished fungal genome." MBio 6(4): e00936–00915.

Goodwin, S., J. Gurtowski, S. Ethe-Sayers, P. Deshpande, M. C. Schatz and W. R. McCombie (2015). "Oxford Nanopore sequencing, hybrid error correction, and de novo assembly of a eukaryotic genome." Genome research 25(11): 1750–1756.

Graziani, G., S. Schiavo, M. A. Nicolai, S. Buono, V. Fogliano, G. Pinto and A. Pollio (2013). "Microalgae as human food: chemical and nutritional characteristics of the thermo-acidophilic microalga Galdieria sulphuraria." Food & function 4(1): 144–152.

Gross, W., J. Küver, G. Tischendorf, N. Bouchaala and W. Büsch (1998). "Cryptoendolithic growth of the red alga Galdieria sulphuraria in volcanic areas.” European Journal of Phycology 33(1): 25–31.

Gross, W. and C. Schnarrenberger (1995). "Heterotrophic growth of two strains of the acido-thermophilic red alga Galdieria sulphuraria." Plant and Cell Physiology 36(4): 633–638.

Henkanatte–Gedera, S., T. Selvaratnam, N. Caskan, N. Nirmalakhandan, W. Van Voorhies and P. J. Lammers (2015). "Algal-based, single-step treatment of urban wastewaters." Bioresource technology 189: 273–278.

Henkanatte–Gedera, S., T. Selvaratnam, M. Karbakhshravari, M. Myint, N. Nirmalakhandan, W. Van Voorhies and P. J. Lammers (2016). "Removal of dissolved organic carbon and nutrients from urban wastewaters by Galdieria sulphuraria: Laboratory to field scale demonstration." Algal Research.

Hsieh, C. J., S. H. Zhan, Y. Lin, S. L. Tang and S. L. Liu (2015). "Analysis of rbcL sequences reveals the global biodiversity, community structure, and biogeographical pattern of thermoacidophilic red algae (Cyanidiales)." Journal of phycology 51(4): 682–694.

Jain, M., I. T. Fiddes, K. H. Miga, H. E. Olsen, B. Paten and M. Akeson (2015). "Improved data analysis for the MinION nanopore sequencer." Nature methods 12(4): 351–356.

Ju, X., K. Igarashi, S.-i. Miyashita, H. Mitsuhashi, K. Inagaki, S.-i. Fujii, H. Sawada, T. Kuwabara and A. Minoda (2016). "Effective and selective recovery of gold and palladium ions from metal wastewater using a sulfothermophilic red alga, Galdieria sulphuraria." Bioresource technology 211: 759–764.

Koch, W. (1964). "Verzeichnis der Sammlung von Algenkulturen am Pflanzenphysiologischen Institut der Universität Göttingen." Archives of Microbiology 47(4): 402–432.

Koren, S., M. C. Schatz, B. P. Walenz, J. Martin, J. T. Howard, G. Ganapathy, Z. Wang, D. A. Rasko, W. R. McCombie and E. D. Jarvis (2012). "Hybrid error correction and de novo assembly of single-molecule sequencing reads.” Nature biotechnology 30(7): 693–700.

Li, H. (2016). “Minimap and miniasm: fast mapping and de novo assembly for noisy long sequences.” Bioinformatics: btw152.

Linka, M., A. Jamai and A. P. Weber (2008). “Functional characterization of the plastidic phosphate translocator gene family from the thermo-acidophilic red alga Galdieria sulphuraria reveals specific adaptations of primary carbon partitioning in green plants and red algae." Plant physiology 148(3): 1487–1496.

Loman, N. J., J. Quick and J. T. Simpson (2015). "A complete bacterial genome assembled de novo using only nanopore sequencing data." Nature methods.

López–Archilla, A. I., I. Marín and R. Amils (2001). "Microbial community composition and ecology of an acidic aquatic environment: the Tinto River, Spain." Microbial ecology 41(1): 20–35.

Merola, A., R. Castaldo, P. D. Luca, R. Gambardella, A. Musacchio and R. Taddei (1981). "Revision of Cyanidium caldarium. Three species of acidophilic algae." Plant Biosystem 115(4–5): 189–195.

Milne, I., D. Lindner, M. Bayer, D. Husmeier, G. McGuire, D. F. Marshall and F. Wright (2009). "TOPALi v2: a rich graphical interface for evolutionary analyses of multiple alignments on HPC clusters and multi-core desktops." Bioinformatics 25(1): 126–127.

Minoda, A., H. Sawada, S. Suzuki, S.-i. Miyashita, K. Inagaki, T. Yamamoto and M. Tsuzuki (2015). "Recovery of rare earth elements from the sulfothermophilic red alga Galdieria sulphuraria using aqueous acid." Applied microbiology and biotechnology 99(3): 1513–1519

Moreira, D., A.–I. López-Archilla, R. Amils and I. Marín (1994). "Characterization of two new thermoacidophilic microalgae: genome organization and comparison with Galdieria sulphuraria." FEMS microbiology letters 122(1–2): 109–114.

Oesterhelt, C., S. Klocke, S. Holtgrefe, V. Linke, A. P. Weber and R. Scheibe (2007). "Redox regulation of chloroplast enzymes in Galdieria sulphuraria in view of eukaryotic evolution." Plant and Cell Physiology 48(9): 1359–1373.

Parker, H. L., E. L. Rylott, A. J. Hunt, J. R. Dodson, A. F. Taylor, N. C. Bruce and J. H. Clark (2014). "Supported palladium nanoparticles synthesized by living plants as a catalyst for Suzuki-Miyaura Reactions." PLoS One 9(1): e87192.

Pinto, G. (2007). Cyanidiophyceae. Algae and Cyanobacteria in Extreme Environments, Springer: 387–397.

Qiu, H., D. C. Price, A. P. Weber, V. Reeb, E. C. Yang, J. M. Lee, S. Y. Kim, H. S. Yoon and D. Bhattacharya (2013). "Adaptation through horizontal gene transfer in the cryptoendolithic red alga Galdieria phlegrea.” Current Biology 23(19): R865–R866.

Schmidt, R. A., M. G. Wiebe and N. T. Eriksen (2005). "Heterotrophic high cell‐density fed‐batch cultures of the phycocyanin‐producing red alga Galdieria sulphuraria." Biotechnology and bioengineering 90(1): 77–84.

Schönknecht, G., W.–H. Chen, C. M. Ternes, G. G. Barbier, R. P. Shrestha, M. Stanke, A. Bräutigam, B. J. Baker, J. F. Banfield and R. M. Garavito (2013). "Gene transfer from bacteria and archaea facilitated evolution of an extremophilic eukaryote." Science 339(6124): 1207–1210.

Schönknecht, G., A. P. Weber and M. J. Lercher (2014). "Horizontal gene acquisitions by eukaryotes as drivers of adaptive evolution." Bioessays 36(1): 9–20.

Selvaratnam, T., A. Pegallapati, F. Montelya, G. Rodriguez, N. Nirmalakhandan, P. J. Lammers and W. Van Voorhies (2015). "Feasibility of algal systems for sustainable wastewater treatment." Renewable Energy 82: 71–76.

Selvaratnam, T., A. Pegallapati, F. Montelya, G. Rodriguez, N. Nirmalakhandan, W. Van Voorhies and P. Lammers (2014). "Evaluation of a thermo-tolerant acidophilic alga, Galdieria sulphuraria, for nutrient removal from urban wastewaters." Bioresource technology 156:395–399.

Skorupa, D., V. Reeb, R. Castenholz, D. Bhattacharya and T. McDermott (2013). “Cyanidiales diversity in Yellowstone National park." Letters in applied microbiology 57(5): 459–466.

Toplin, J., T. Norris, C. Lehr, T. McDermott and R. Castenholz (2008). "Biogeographic and phylogenetic diversity of thermoacidophilic cyanidiales in Yellowstone National Park, Japan, and New Zealand." Applied and environmental microbiology 74(9): 2822–2833.

Weber, A., C. Oesterhelt, W. Gross, A. Bräutigam, L. Imboden, I. Krassovskaya, N. Linka, J. Truchina, J. Schneidereit and H. Voll (2004). "EST-analysis of the thermo-acidophilic red microalga Galdieriasulphuraria reveals potential for lipid A biosynthesis and unveils the pathway of carbon export from rhodoplasts." Plant molecular biology 55(1): 17–32.

Willing, E.–M., V. Rawat, T. Mandáková, F. Maumus, G. V. James, K. J. Nordström, C. Becker, N. Warthmann, C. Chica and B. Szarzynska (2015). "Genome expansion of Arabis alpina linked with retrotransposition and reduced symmetric DNA methylation." Nature Plants 1(2).

Yoon, H., C. Ciniglia, M. Wu, J. M. Comeron, G. Pinto, A. Pollio and D. Bhattacharya (2006). “Establishment of endolithic populations of extremophilic Cyanidiales (Rhodophyta)." BMC evolutionary biology 6(1): 1.

